# Predicting Biological Age and Clinical Biomarkers from DNA Methylation Profiles of Cheek Mucosa

**DOI:** 10.64898/2026.05.12.724485

**Authors:** Tatsuma Shoji, Yui Tomo, Ryo Nakaki

## Abstract

**Background:** DNA methylation-based biomarkers have been widely used to predict biological age; however, most blood-derived data have been used in most existing models, and whether cheek mucosa can serve as an alternative indicator for methylation-based estimation of aging-related and clinical phenotypes is unclear.

**Methods:** DNA methylation profiles from cheek mucosa and whole blood of 186 Japanese adults were analyzed using Illumina Infinium Methylation Screening Array (MSA). Models were constructed to predict chronological age, phenotypic age, and clinical laboratory biomarkers from cheek mucosa- and blood-derived methylation data. In addition to applying the ordinary elastic net method, a two-stage residual learning method incorporating existing blood-based epigenetic clocks was applied for more accurate prediction of biological age. Sex-stratified analyses and comparisons of selected CpG features across sexes and tissues were performed.

**Results:** Cheek mucosa-derived MSA methylation data enabled accurate prediction of chronological age (R = 0.965) and phenotypic age (R = 0.964) using the two-stage method. The performance gain achieved by the two-stage approach was greater for phenotypic age than for chronological age. Multiple clinical laboratory biomarkers could be predicted using cheek mucosa-derived methylation data, particularly after sex stratification, including inflammatory, metabolic, thyroid-related, and sex hormone-related markers. Most biomarkers that could be predicted using blood-derived methylation data were also predicted using cheek mucosa-derived methylation data. However, the CpG sites selected for prediction showed minimal overlap across sexes and tissues despite overlap in the corresponding predictable phenotypes.

**Conclusions:** Cheek mucosa-derived DNA methylation profiles measured using the MSA can predict chronological age, phenotypic age, and multiple clinically relevant laboratory biomarkers, supporting the utility of cheek mucosa as a less invasive alternative for methylation-based assessment of biological aging and systemic physiological state.

## Introduction

DNA methylation-based biomarkers are crucial for quantifying biological aging in a clinically interpretable manner [1–3]. Unlike chronological age, biological age intends to capture inter-individual variation in physiological decline, disease vulnerability, and mortality risk, making it potentially useful for risk stratification and longitudinal health assessment [2, 4–6]. Early epigenetic clocks, including the Horvath and Hannum models, demonstrated that age-associated methylation patterns could predict chronological age with high accuracy across human samples, effectively predicting the age of a typical healthy individual corresponding to a given DNA methylation profile, and established DNA methylation as a reliable molecular biomarker of aging [3, 7]. More recently, epigenetic clocks or second-generation clocks such as PhenoAge and GrimAge have improved the clinical relevance of this approach by incorporating mortality- and health-related phenotypes rather than chronological time alone [2, 8]. These advancements support the use of DNA methylation profiling as a scalable approach for clinically relevant aging assessment and for the development of molecular biomarkers for better accessibility in monitoring systemic health states.

Despite these advances, several limitations constrain the clinical applicability of DNA methylation-based aging biomarkers. Most established epigenetic clocks have been developed and validated primarily using blood-derived DNA, despite the invasive nature of blood collection and potential limitations of repeated or large-scale implementation in screening and longitudinal monitoring settings, particularly due to the procedural burden and challenges in participant compliance with frequent sampling. DNA methylation profiles are strongly tissue-dependent, and predictive models developed in one tissue may not be directly transferable to another [9, 10]. Simultaneously, buccal and oral epithelial samples are attractive alternatives for translational applications because they can be collected non-invasively, repeatedly, and with less burden on participants. Indeed, buccal methylation data can support accurate age prediction and may capture biologically relevant aging-related variation, including tissue-specific age predictors such as PedBE, and more recent buccal aging models have been developed for adult populations [11–14]. However, the extent to which buccal or cheek mucosa-derived methylation profiles can recover clinically relevant aging phenotypes and blood-associated physiological information remains insufficiently characterized, particularly when using a methylation screening array (MSA) and in non-European populations.

Beyond age prediction, DNA methylation profiles may capture information related to systemic physiological states and circulating clinical biomarkers. DNA methylation signatures can be used to predict blood-based traits, including inflammatory markers, immune-related measures, lipid-related indices, and hormone-associated phenotypes, suggesting that methylation reflects integrative biological signals shaped by both intrinsic regulation and environmental exposure [2, 15]. Such DNA methylation-derived biomarker proxies are associated with morbidity, functional decline, and mortality risk, highlighting their potential applicability for epidemiological and translational research. We recently showed that multiple clinical laboratory biomarkers could be predicted from blood-derived methylation data generated using the Illumina Infinium MSA [16], supporting the feasibility of biomarker prediction even on a lower-density, cost-efficient platform. However, whether a similar physiological inference is feasible using non-blood tissues, particularly less invasive approaches such as sampling cheek mucosa, remains unclear. MSA is enriched for CpG sites relevant to aging, inflammation, metabolism, and chronic disease, making it a promising platform for extending methylation-based phenotyping beyond age prediction. Nevertheless, any evaluation of whether clinically relevant blood biomarkers can be predicted from cheek mucosa-derived MSA methylation data, and how such predictions compare with those obtained from blood-derived MSA data, is lacking.

Recently, we developed a Japanese population-based methylation predictor of phenotypic age, a composite measure of biological age derived from clinical biomarkers predictive of mortality risk [2], using Infinium EPIC v2 data [16], and subsequently J-PhenoAge [17], an MSA-based model for estimating phenotypic age from blood-derived methylation profiles. These studies suggested that population-specific and platform-adapted methylation models may improve the utility of biological age estimation in Japanese cohorts. In this study, we investigated whether DNA methylation profiles obtained from the cheek mucosa using the Illumina Infinium MSA could be used to predict clinically relevant aging phenotypes in a Japanese cohort. Specifically, we evaluated whether cheek mucosa-derived MSA data could predict chronological age and phenotypic age, and whether a residual learning approach could improve predictive performance by leveraging information from established blood-based epigenetic clocks. We compared cheek mucosa-derived predictions with blood-derived methylation-based ones and examined whether individual clinical laboratory biomarkers could be predicted from cheek mucosa and blood MSA data. We assessed the overlap and divergence of CpG features selected across tissues and sexes to better characterize the biological and technical specificity underlying these prediction models. By integrating tissue comparison, biomarker prediction, and feature-level analyses, this study aimed to evaluate the feasibility and limitations of using a less invasive oral sampling approach as a clinically accessible method to assess methylation-based aging and physiological state.

## Material & Methods

### Study Participants and Sample Collection

Whole blood and cheek mucosa samples were collected from healthy adult volunteers under an approved observational study protocol. A total of 186 unique participants were enrolled, and paired cheek mucosa and whole blood samples were obtained at Y’s Science Clinic Hiroo (Minato-ku, Tokyo, Japan) as part of the project titled “Evaluation of Biological Age Based on DNA Methylation and Its Clinical Significance,” reviewed by the Institutional Review Board of Shiba Palace Clinic. Written informed consent was obtained from all participants before sample collection. Genomic DNA was extracted from whole blood using the Maxwell RSC Blood DNA Kit (Promega, Madison, WI, USA) following the manufacturer’s instructions. Genomic DNA from cheek mucosa samples was extracted using Maxwell RSV Stabilized Saliva DNA Kit (Promega, Madison, WI, USA) following the manufacturer’s instructions. All procedures were conducted in accordance with the Ethical Guidelines for Medical and Biological Research Involving Human Subjects issued by the Japanese government.

### Clinical Laboratory Measurements

Fifty-nine clinical laboratory markers were measured in whole blood samples collected at the time of DNA extraction, following the protocol described previously in Shoji et al [16]. Briefly, the measured variables included standard biochemical markers, hematological indices, hormone-related parameters, inflammatory markers, lipid-related measurements, and serum protein fractionation profiles. The assessed markers included: alkaline phosphatase based on the International Federation of Clinical Chemistry and Laboratory Medicine (ALP/IFCC), alanine aminotransferase (ALT), aspartate aminotransferase (AST), quantitative C-reactive protein (CRP), calcium (Ca), chloride (Cl), dehydroepiandrosterone sulfate (DHEA-S), follicle-stimulating hormone (FSH), serum iron, hematocrit, high-density lipoprotein (HDL) cholesterol, hemoglobin (Hb), hemoglobin A1c (HbA1c), potassium (K), lactate dehydrogenase based on IFCC (LD/IFCC), low-density lipoprotein (LDL) cholesterol, luteinizing hormone (LH), mean corpuscular hemoglobin (MCH), mean corpuscular hemoglobin concentration (MCHC), mean corpuscular volume (MCV), magnesium (Mg), sodium (Na), N-terminal pro-brain natriuretic peptide (NT-proBNP), phosphorus (P), red blood cell (RBC) count, red cell distribution width-coefficient of variation (RDW-CV), thyroid-stimulating hormone (TSH), white blood cell (WBC) count, γ-glutamyl transpeptidase (γ-GTP), albumin, insulin, estradiol, creatinine, total testosterone, free triiodothyronine (free T3), free thyroxine (free T4), progesterone, triglycerides, blood urea nitrogen (BUN), uric acid, differential leukocyte counts (lymphocytes, monocytes, neutrophils, basophils, and eosinophils), total cholesterol, total bilirubin, total protein, α1-globulin, α2-globulin, β-globulin, and γ-globulin, albumin/globulin (A/G) ratio, albumin fraction, platelet count, serum amylase, blood glucose, free testosterone, and high-sensitivity prostate-specific antigen (PSA).

### Outlier Detection and Missing-Value Imputation

Outlier detection and missing-value imputation were performed as described in Shoji et al [16]. Outlier detection was conducted independently for each clinical laboratory marker using a conditional residual approach. For each marker, a Huber regression model with epsilon = 1.35 was fitted using the remaining 58 laboratory markers as predictors, and residuals were standardized using the median absolute deviation. Values with a standardized conditional residual ≥ 6 were flagged as potential outliers. Samples with statistically significant Mahalanobis distances in the standardized feature space, as determined by a chi-square test at a significance level of 0.0001, were considered potential multivariate outliers. No individual measurements or samples met the exclusion criteria. Missing values were imputed using a multivariate single-imputation strategy based on the ExtraTrees algorithm [18]. The proportions of missing values for each laboratory marker are summarized in Additional file 1, and the imputed values are provided in Additional file 2. Outlier detection and missing-value imputation were performed using Scikit-learn (version 1.7.1) in Python (version 3.12.3).

### Calculating the Phenotypic Age

Phenotypic age was calculated following the procedure described in the original PhenoAge study [2]. This approach calculated a mortality-associated biological age based on nine clinical biomarkers. Albumin values were calculated from total protein and albumin fractions obtained through serum protein fractionation, while the remaining biomarkers were converted to respective units. The nine biomarkers included: albumin (g/L), creatinine (µmol/L), blood glucose (mmol/L), C-reactive protein (mg/L), lymphocyte percentage (%), mean corpuscular volume (MCV, fL), red cell distribution width-coefficient of variation (RDW-CV, %), alkaline phosphatase (ALP, U/L), and white blood cell (WBC) count (10³ cells/µL). After harmonizing measurement units, these variables were incorporated into the published mortality score equation. Specifically, the mortality score (MS) was calculated as:

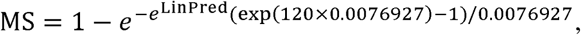

where

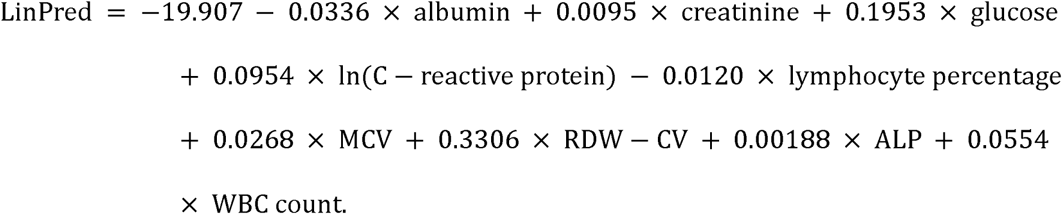

Phenotypic age was calculated by transforming this mortality score according to the following equation:

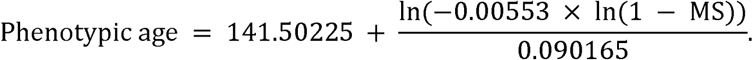

All calculations were performed using Python (version 3.12.3).

### DNA Methylation Profiling and Preprocessing

Genome-wide DNA methylation profiling was performed in a manner similar to that described by Shoji et al. [16]. Genomic DNA derived from both cheek mucosa and whole blood samples was analyzed using the Illumina Infinium MSA, whereas genomic DNA derived from whole blood samples was additionally analyzed using Illumina Infinium HumanMethylationEPIC v2 BeadChip (EPICv2). Rhelixa, Inc. (Chuo-ku, Tokyo, Japan) processed and scanned the array. Following bisulfite conversion according to the manufacturer’s recommended protocol, the converted DNA was amplified, fragmented, hybridized to the arrays, and scanned using an Illumina iScan System. Raw IDAT files were processed in R (version 4.4.2) using the SeSAMe pipeline (version 1.24.0) [19, 20], including background correction with the normal-exponential out-of-band (noob) method and probe-level quality filtering using the pOOBAH detection mask. DNA methylation beta values were calculated as the ratio of methylated signal intensity to the sum of methylated and unmethylated signal intensities [21]. Before downstream analyses, probes that failed detection thresholds, those overlapping with single-nucleotide polymorphisms likely to affect hybridization, those exhibiting cross-reactive mapping to multiple genomic loci, or those located on sex chromosomes were excluded. Raw IDAT files and processed beta value matrices are available upon reasonable request.

### Development of Cheek Mucosa-Derived MSA-Based Age Prediction Models

To develop cheek mucosa-derived MSA-based prediction models for chronological age and phenotypic age, linear regression models were trained using CpG methylation beta values derived from cheek mucosa MSA data as predictors and chronological age or Phenotypic Age as the target variable. The Elastic Net method was used for model training [20]. For each model, all predictors were standardized before training. The dataset comprising 186 participants was randomly divided such that 80% of the samples (n=149) were used for hyperparameter tuning with fivefold cross-validation, whereas the remaining 20% were reserved as a held-out test set. After identifying the optimal hyperparameters from candidates, the final model performance was evaluated in the held-out test set by calculating the Pearson correlation coefficient, mean absolute error (MAE), and root mean squared error (RMSE) between predicted and observed values. The CpG sites selected in the final models and their corresponding regression coefficients are summarized in Additional file 3. All model training and visualization were performed using Scikit-learn (version 1.7.1) and Matplotlib (version 3.10.6) in Python (version 3.12.3).

### Computation of Existing Epigenetic Clocks

For comparison with the cheek mucosa-derived MSA-based prediction models, previously published epigenetic clocks were computed using blood-derived DNA methylation data. The Horvath, Hannum, and PhenoAge clocks were calculated using blood-derived EPIC v2 methylation data using the CpG sets and regression coefficients reported in their original publications [1, 7]. Five methylation-based age-related indicators, Horvath, Hannum, PhenoAge, GrimAgeV2, and J-PhenoAge, were calculated from blood-derived MSA methylation data according to previously described procedures [1, 2, 7, 16, 22]. Briefly, each indicator was computed using the corresponding published regression coefficients. When CpG sites required for a given clock were not available on the MSA platform, their coefficients were set to zero for score calculation.

### Model Development using the Two-Stage Residual Learning Approach

A two-stage residual learning approach was implemented to evaluate whether predictive performance could be improved by incorporating information from existing epigenetic clocks, following the approach described by Shoji et al. [16]. Specifically, this approach comprises two steps: 1. computing residuals from a first-stage linear regression model where chronological age or phenotypic age is the target variable and the five epigenetic ages computed by the existing epigenetic clocks as the predictors; 2. specifying the residuals as the target variable for a second-stage model and MSA-measured CpGs as the predictors. This approach can be viewed as a form of transfer learning, where information captured by established epigenetic clocks is refined using a new set of methylation features [21, 23]. In our analysis, in the first stage, five epigenetic clocks (Horvath, Hannum, PhenoAge, GrimAgeV2, and J-PhenoAge) were computed and standardized for predictors in a linear regression model trained using *l_2_* regularization (Ridge regression) to estimate the target variable. In the second stage, residuals obtained by subtracting the first-stage predicted values from the observed values were used as target variables, and MSA-derived CpG beta values were used as predictors in linear regression models, and models were trained by the Elastic Net method. For validation, the held-out test set was used to calculate first-stage predicted values and second-stage predicted residuals, which were summed to obtain the final predictions. Model performance was evaluated by calculating the Pearson correlation coefficient, MAE, and RMSE between predicted and observed values. The coefficients used in the first- and second-stage models are summarized in Additional file 4. All calculations and visualizations were performed using Matplotlib (version 3.10.6) and Scikit-learn (version 1.7.1) in Python (version 3.12.3).

### Prediction of Clinical Biomarkers from MSA Data

Predictive models were constructed for all 59 clinical laboratory biomarkers, and for the log-transformed values of six right-skewed variables (γ-GTP, NT-proBNP, triglycerides, AST, ALT, and quantitative CRP), following the modeling strategy described by Shoji et al. [16]. Two modeling approaches were implemented using either blood-derived or cheek mucosa-derived MSA methylation data to assess whether individual biomarkers could be predicted from methylation-derived features. In Model 1, linear regression models were constructed using CpG methylation beta values as predictors and each clinical biomarker as the target variable. Model training was performed using the Elastic Net method on 80% of the dataset (n = 149), with fivefold cross-validation used for hyperparameter tuning, while the remaining 20% (n = 37) was reserved as a held-out test set. Predictive performance was assessed in the held-out test set by calculating the Pearson correlation coefficient, MAE, and RMSE between observed and predicted values. Partial correlation coefficients controlling for sex were calculated to account for potential confounding by sex. In Model 2, participants were stratified by sex, and separate models were trained for males and females using CpG methylation beta values and chronological age as predictors. Model performance was evaluated independently within each sex using the held-out test set and summarized using the Pearson correlation coefficient, MAE, and RMSE. The MAE and RMSE values are summarized in Additional File 5. The CpG sites used in the final models for both Model 1 and Model 2, and their corresponding regression coefficients, are summarized in Additional files 6–11. All calculations and visualizations were performed using Scikit-learn (version 1.7.1) and Matplotlib (version 3.10.6) in Python (version 3.12.3).

### Comparative Analysis of Selected CpG Features

Comparative analyses were performed using the CpG features selected in Model 2 to investigate whether the CpG sites selected for biomarker prediction differed by sex and tissue type. First, to evaluate sex-associated differences, the sets of CpG sites selected in the male- and female-specific models were compared for each biomarker in both cheek mucosa-derived and blood-derived MSA models. The degree of overlap between the two sets was quantified using the Jaccard index. M values were calculated for all CpG sites included on the MSA platform, and Welch’s t-tests were performed between males and females to determine whether the selected CpG features were enriched for sex-associated methylation differences. Resulting P values were adjusted for multiple testing using the Benjamini–Hochberg false discovery rate (BH-FDR) method, and CpG sites with an adjusted P value of ≤ 0.1 were defined as sex-associated CpG sites. The proportion of sex-associated CpG sites among those selected in the male- or female-specific prediction models was then calculated. Similarly, to evaluate tissue-associated differences, the CpG sites selected in the cheek mucosa-derived and blood-derived sex-specific models were compared separately for males and females for each biomarker, and overlap was quantified using the Jaccard index. To identify tissue-associated CpG sites, M values were calculated for all CpG sites included on the MSA platform, and Welch’s t-tests were performed between cheek mucosa and blood samples. After applying the BH-FDR method, CpG sites with an adjusted P value of ≤ 0.1 were defined as tissue-associated CpG sites. The proportion of tissue-associated CpG sites among those selected in the cheek mucosa- or blood-derived prediction models was calculated.

## Results

### Prediction of Chronological Age and Phenotypic Age from Cheek Mucosa-Derived MSA Data

We first evaluated whether chronological age and phenotypic age could be predicted from cheek mucosa-derived MSA methylation data and whether predictive performance could be improved using the two-stage residual learning approach. In the held-out test set, the two-stage model predicted chronological age with high accuracy (R = 0.965, MAE = 2.602, RMSE = 3.311)(Fig. 1A) and showed strong performance for predicting phenotypic age (R = 0.964, MAE = 3.405, RMSE = 4.217)(Fig. 1B). By comparison, the corresponding one-stage models showed lower predictive performance for both chronological age (R = 0.947, MAE = 3.277, RMSE = 4.118)(Fig. 1C) and phenotypic age (R = 0.854, MAE = 6.214, RMSE = 8.432)(Fig. 1D). The difference between the one-stage and two-stage models was more pronounced for phenotypic age than for chronological age.

**Figure 1.**
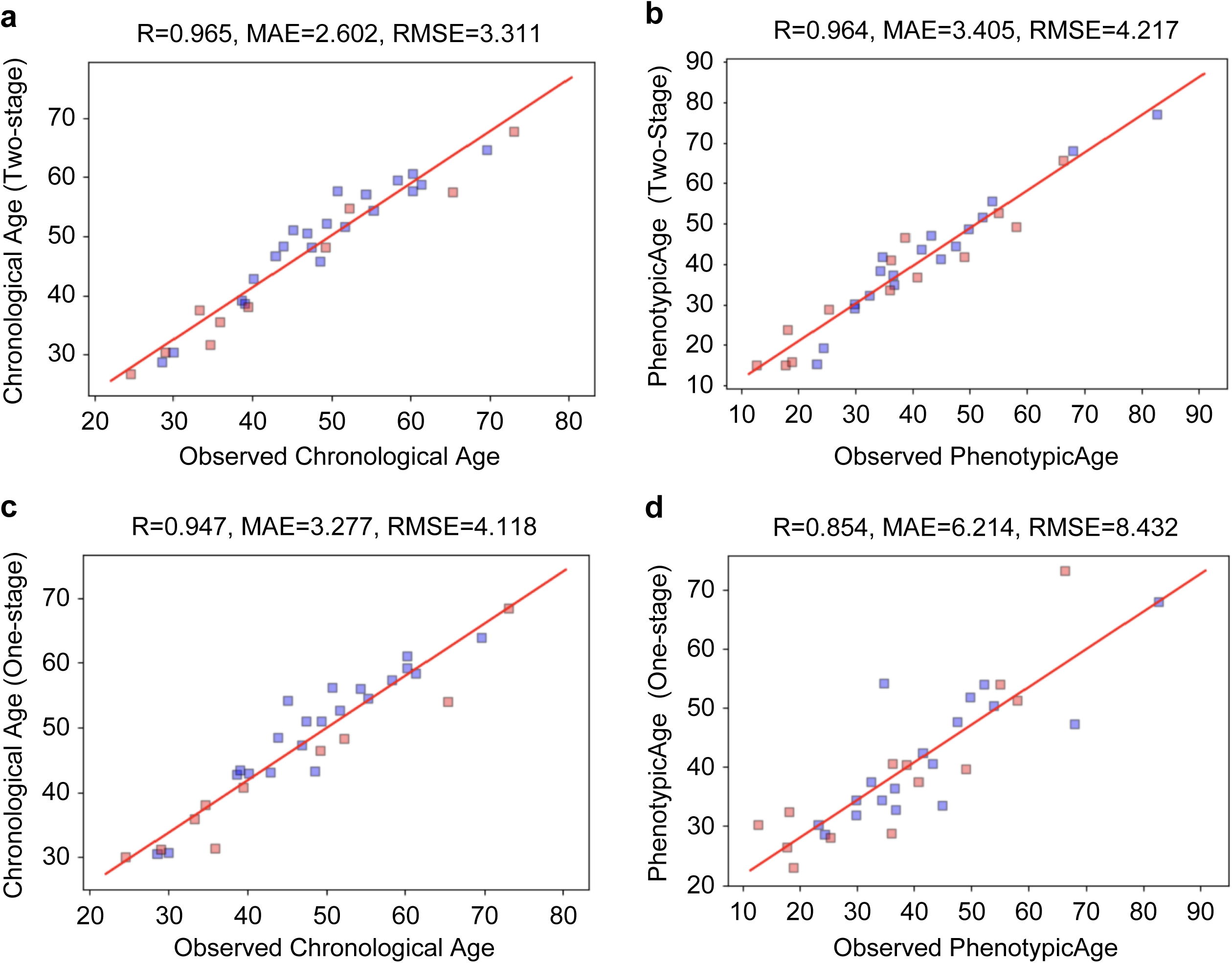
Comparison of observed and predicted chronological age and phenotypic age. Scatter plots showing observed versus predicted values for chronological age and phenotypic age. Predictions were based on the cheek mucosa-derived MSA-based two-stage chronological age model (A), the cheek mucosa-derived MSA-based two-stage phenotypic age model (B), the cheek mucosa-derived MSA-based one-stage chronological age model (C), and the cheek mucosa-derived MSA-based one-stage phenotypic age model (D). Blue dots indicate male participants, and red dots indicate female participants. The correlation coefficient (R), mean absolute error (MAE), and root mean squared error (RMSE) are shown at the top of each panel.

### Comparison with Blood-based Epigenetic Clocks

We next examined how the cheek mucosa-derived MSA-based prediction models related to established blood-based epigenetic clocks. For chronological age-related measures, both the one-stage and two-stage cheek mucosa-derived models showed strong correlations with the blood-derived Horvath (R = 0.944, MAE = 4.543, RMSE = 5.699 for the one-stage model and R = 0.911, MAE = 5.315, RMSE = 6.631 for the two-stage model)(Fig. 2A, B) and Hannum clocks (R = 0.953, MAE = 7.908, RMSE = 8.674 for the one-stage model and R = 0.930, MAE = 4.671, RMSE = 5.551 for the two-stage model) (Fig. 2C, D). In contrast, for phenotypic age-related measures, the corresponding cheek mucosa-derived models showed comparatively lower correlations with the blood-derived PhenoAge clock (R = 0.894, MAE = 7.224, RMSE = 8.779 for one-stage model and R = 0.841, MAE = 6.211, RMSE = 8.006 for two-stage model) (Fig. 2E, F). Overall, the cheek mucosa-derived models were more strongly aligned with blood-based chronological age indicators than with the blood-based phenotypic age-related measure.

**Figure 2.**
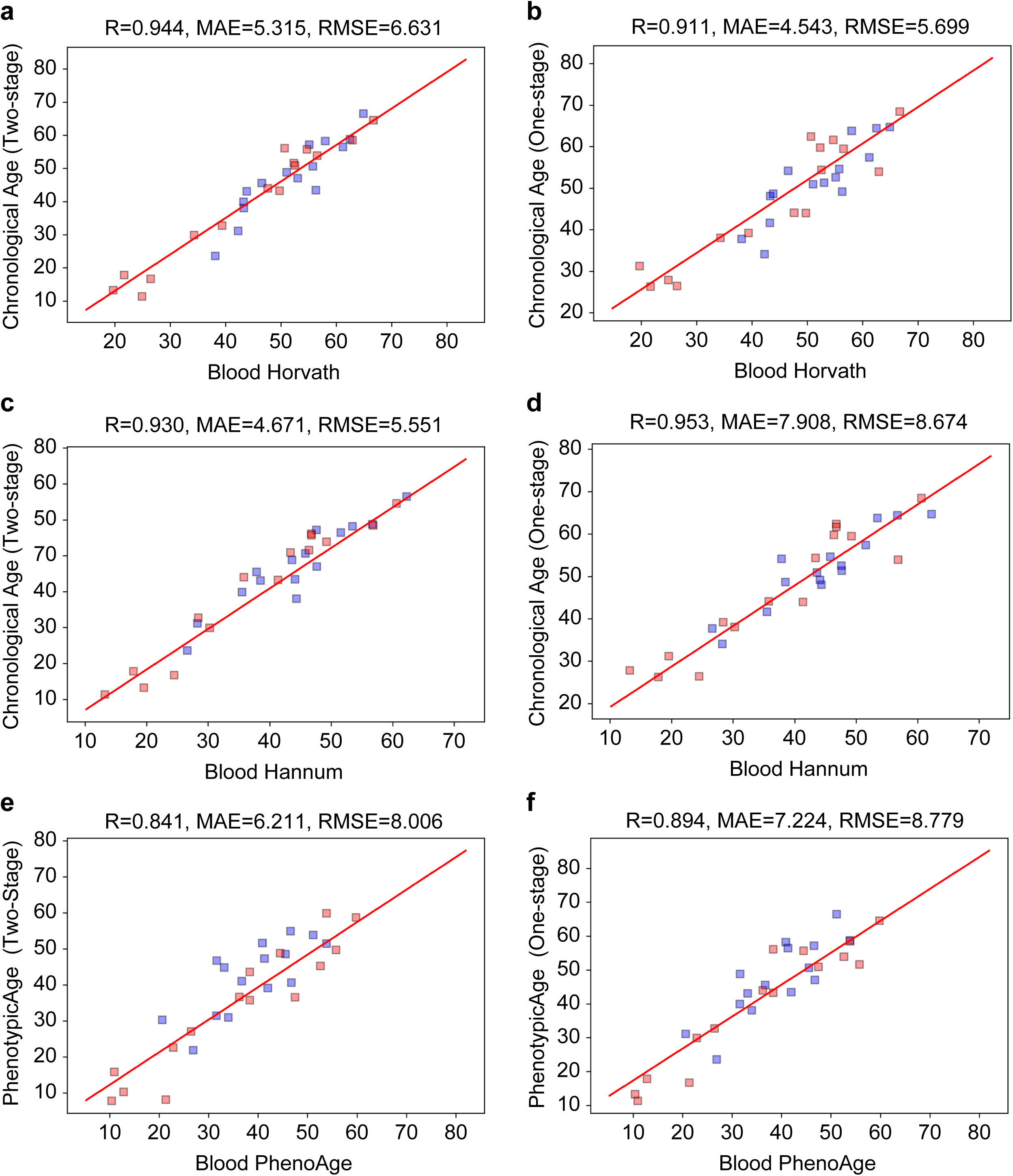
Comparison of cheek mucosa-derived MSA-based age predictions with blood-derived epigenetic clocks. Scatter plots showing the relationships between cheek mucosa-derived MSA-based age predictions and blood-derived epigenetic clock predictions. Panels show comparisons between the cheek mucosa-derived MSA-based two-stage chronological age model and the blood-derived Horvath clock (A), the cheek mucosa-derived MSA-based one-stage chronological age model and the blood-derived Horvath clock (B), the cheek mucosa-derived MSA-based two-stage chronological age model and the blood-derived Hannum clock (C), the cheek mucosa-derived MSA-based one-stage chronological age model and the blood-derived Hannum clock (D), the cheek mucosa-derived MSA-based two-stage phenotypic age model and the blood-derived PhenoAge clock (E), and the cheek mucosa-derived MSA-based one-stage phenotypic age model and the blood-derived PhenoAge clock (F). Blue dots indicate male participants, and red dots indicate female participants. The correlation coefficient (R), mean absolute error (MAE), and root mean squared error (RMSE) are shown at the top of each panel.

### Prediction of Clinical Biomarkers from Cheek Mucosa-Derived MSA Data

We next assessed whether individual clinical laboratory biomarkers could be predicted from cheek mucosa-derived MSA methylation data. Across all participants, several biomarkers showed relatively high predictive performance in the test set, including creatinine and free testosterone (Fig. 3, R [All] > 0.6). However, because some biomarkers appeared to cluster by sex, we evaluated prediction performance using partial correlation coefficients adjusted for sex. After controlling for sex, the associations between observed and predicted values were attenuated for several biomarkers, including creatinine and hemoglobin (Fig. 3, Partial R). Under this criterion, free testosterone was the only biomarker that remained comparatively well predicted from cheek mucosa-derived methylation data (R > 0.6 and partial R > 0.3).

**Figure 3.**
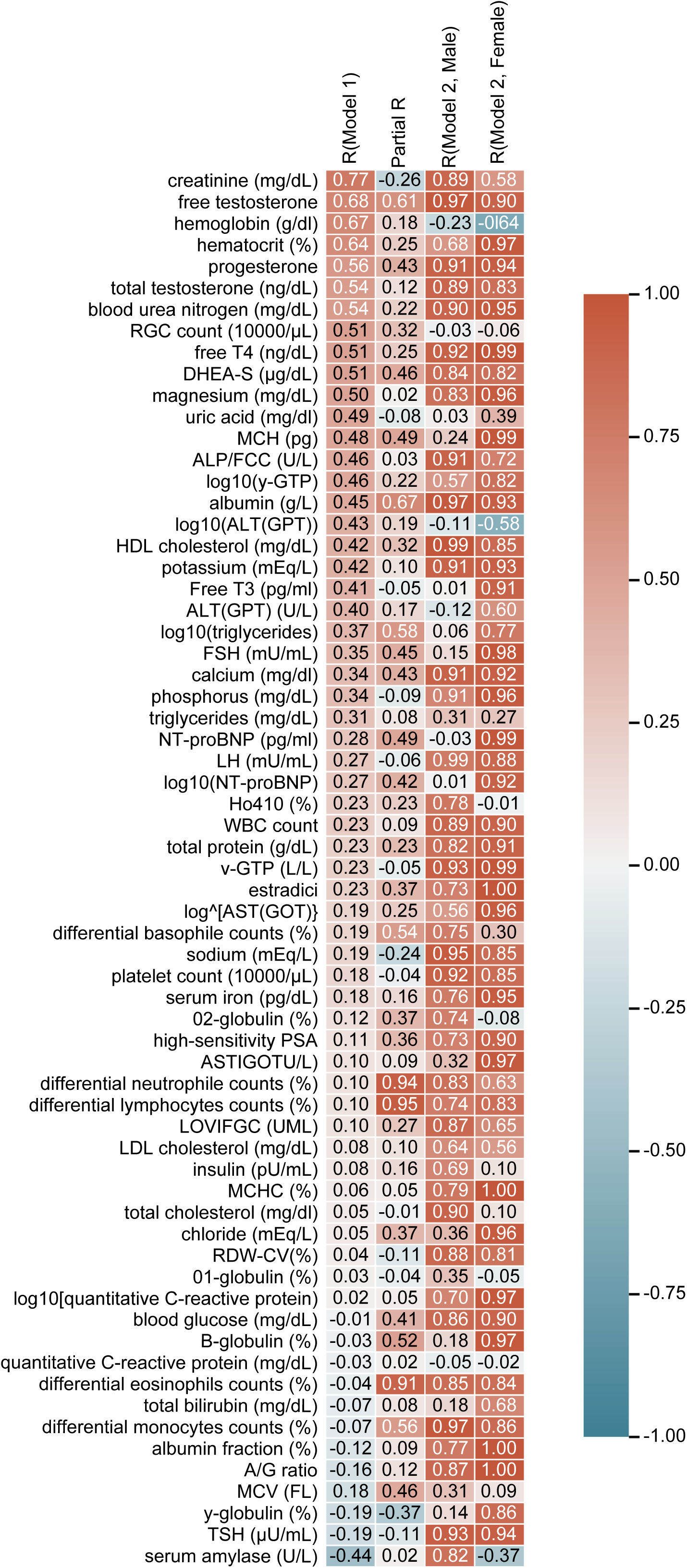
Prediction performance for clinical biomarkers based on cheek mucosa-derived MSA methylation data. Heatmap showing the correlation coefficients between observed and predicted values for all 59 clinical biomarkers. Model 1 represents the prediction performance across all participants and is shown as the Pearson correlation coefficient (R, Model 1). Partial R indicates the sex-adjusted partial correlation coefficient. Model 2 represents the sex-stratified prediction performance and is shown separately for males (R, Model 2 Male) and females (R, Model 2 Female).

### Sex-Stratified Prediction of Clinical Biomarkers from Cheek Mucosa-Derived MSA Data

We next constructed sex-stratified models using cheek mucosa-derived MSA methylation data to evaluate the impact of sex on biomarker prediction performance. Stratification by sex generally improved predictive performance across multiple biomarkers (Fig. 3, Fig. 4A, B). In both males and females, relatively high prediction performance (R > 0.6) was observed for a broad set of biomarkers, including phosphorus, quantitative CRP, estradiol, high-sensitivity PSA, differential lymphocyte counts, LH, serum iron, albumin, Mg, total protein, A/G ratio, blood glucose, differential eosinophil counts, DHEA-S, free T4, γ-GTP, BUN, progesterone, K, Ca, TSH, WBC count, MCHC, platelet count, total testosterone, RDW-CV, HDL cholesterol, free testosterone, total testosterone, differential monocyte counts, Na, ALP/IFCC, LD/IFCC, and differential neutrophil counts. In contrast, hemoglobin, RBC count, uric acid, MCV, and α1-globulin showed relatively low prediction performance in both sexes (R ≤ 0.6) (Fig. 4A). Several biomarkers showed sex-dependent prediction patterns (Fig. 4A). Creatinine, HbA1c, differential basophil counts, α2-globulin, LDL cholesterol, insulin, total cholesterol, and serum amylase were more accurately predicted in males, whereas MCH, free T3, ALT(GPT), triglycerides, FSH, NT-proBNP, AST, Cl, β-globulin, total bilirubin, γ-globulin, and hematocrit were more accurately predicted in females.

**Figure 4.**
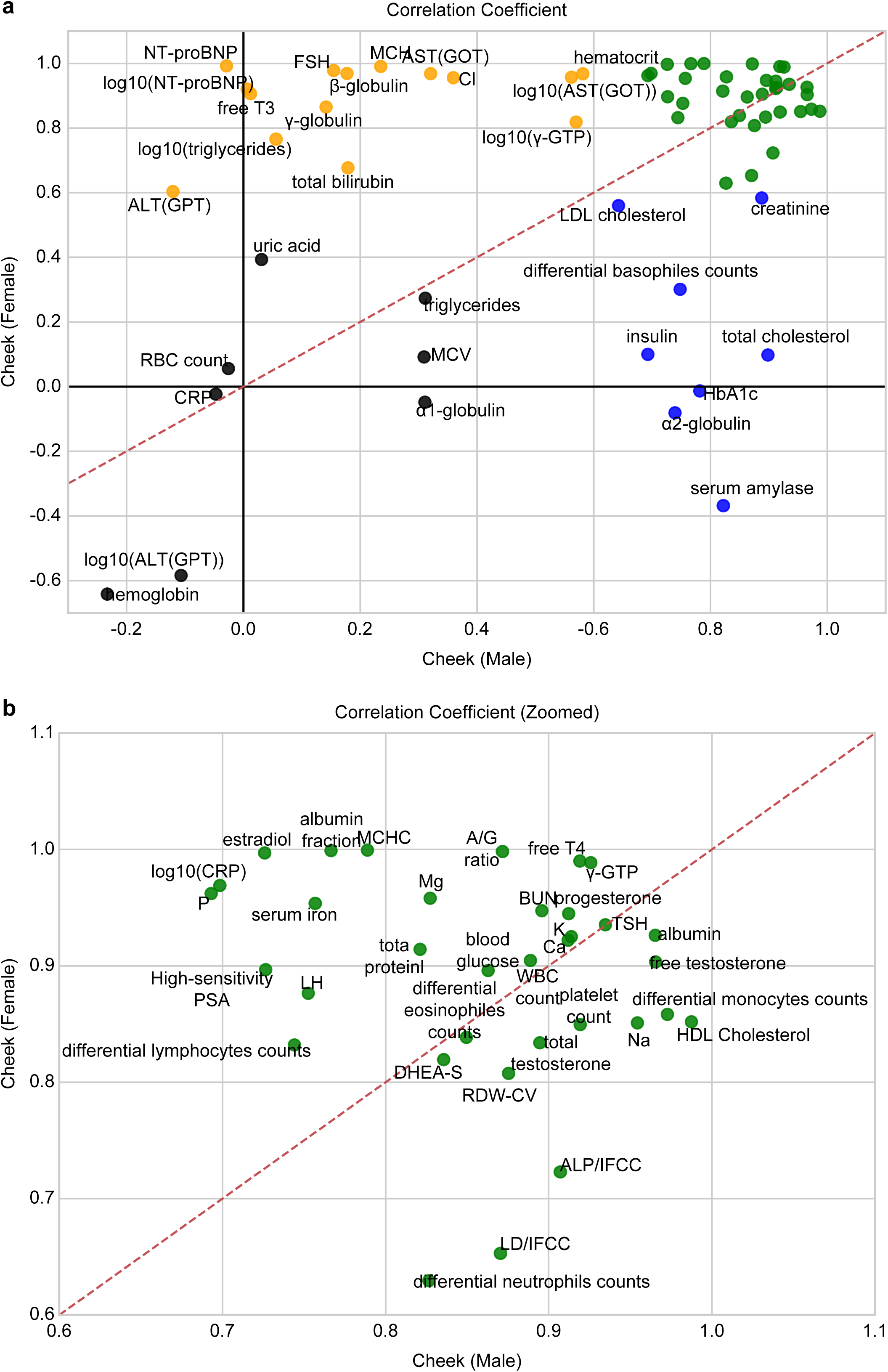
Comparison of biomarker prediction performance between males and females based on cheek mucosa-derived MSA methylation data. Scatter plot showing the comparison of correlation coefficients obtained from Model 2 for all 59 clinical biomarkers between males and females (A). Panel (B) shows an enlarged view of panel (A). Biomarkers with correlation coefficients ≥ 0.6 in both sexes are shown in green, those with correlation coefficients ≥ 0.6 in males only are shown in blue, and those with correlation coefficients ≥ 0.6 in females only are shown in orange. All other biomarkers are shown in black. The red dashed line indicates the line of identity (*y* = *x*).

### Comparison of Selected CpG Features Between Models by Sex

We next examined whether the CpG sites selected in the sex-stratified biomarker prediction models differed between males and females. For most biomarkers, the sets of CpG sites selected in the male and female models showed small overlap, as evidenced by the low Jaccard indices (Fig. 5). To characterize these differences, we assessed whether the selected CpG sites were enriched for sex-associated methylation differences. Across most biomarkers, approximately 20% of the CpG sites selected in the male- or female-specific models overlapped with CpG sites exhibiting significant sex-associated methylation differences, as defined by genome-wide analysis (Fig. 5).

**Figure 5.**
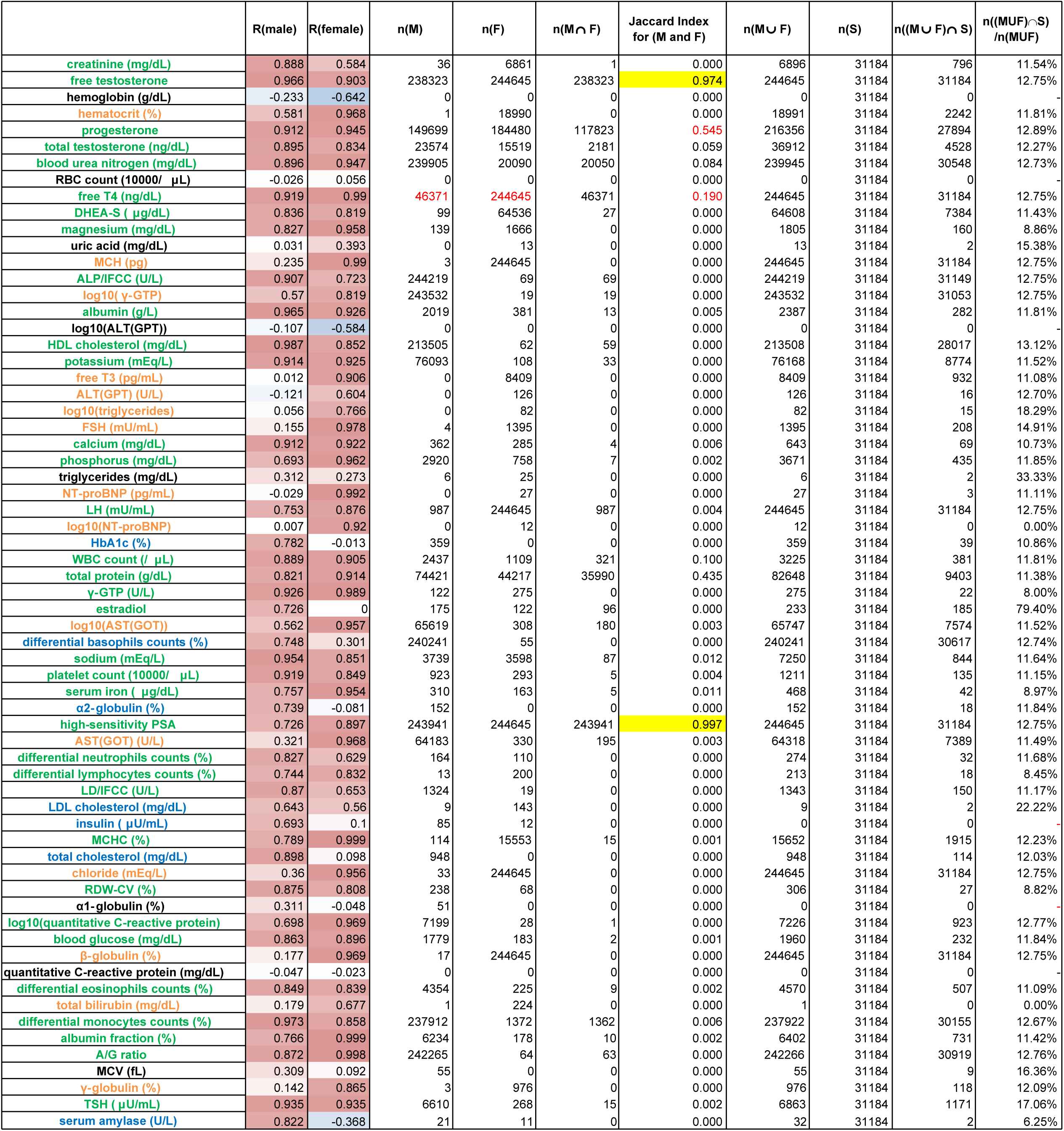
Comparison of CpG features between models by sex based on cheek mucosa-derived DNA methylation and their overlap with sex-associated CpG sites. For all 59 clinical biomarkers, the prediction performance of Model 2 is shown as correlation coefficients for males (R [Male]) and females (R [Female]), with text colors corresponding to Figure 4. The number of CpG sites selected in male (M) and female (F) models, their overlap (n(M ∩ F)), and the Jaccard index are presented, with values ≥ 0.8 highlighted in yellow. When M ∪ F contained no CpG sites, the Jaccard index was set to 0. Overlap with sex-associated CpG sites (S) is evaluated using the number of CpGs in M ∪ F, the number of overlapping CpGs (n((M ∪ F) ∩ S)), and their proportion. When M ∪ F contained no CpG sites, the value is indicated using a hyphen.

### Prediction of Clinical Biomarkers from Blood-Derived MSA Data

We next evaluated whether clinical laboratory biomarkers could be predicted from blood-derived MSA methylation data. Across all participants, several biomarkers showed relatively high predictive performance, including hemoglobin, red blood cell count, creatinine, and free testosterone (Fig. 6, R [All] > 0.6). However, as observed in cheek mucosa-derived models, some of these associations were attenuated after adjusting for sex using partial correlation analysis (Fig. 6, Partial R). Under this criterion, free testosterone remained among the most consistently predicted biomarkers, whereas the predictive performance of several other markers was reduced after accounting for sex. Sex-stratified modeling further improved prediction performance for multiple biomarkers in blood-derived MSA data (Fig. 6, Fig. 7A, B). In both males and females, relatively high prediction performance (R > 0.6) was observed for a broad set of biomarkers, including RBC count, creatinine, phosphorus, estradiol, high-sensitivity PSA, differential lymphocyte counts, LH, albumin, total protein, A/G ratio, ALP(GPT), AST(GOT), differential eosinophil counts, MCHC, DHEA-S, BUN, progesterone, HDL cholesterol, K, Ca, WBC count, platelet count, β-globulin, total testosterone, RDW-CV, NT-proBNP, free testosterone, total testosterone, differential monocyte counts, differential basophil counts, Na, Cl, FSH, LDL cholesterol, ALP/IFCC, LD/IFCC, and differential neutrophil counts. Several biomarkers also showed sex-dependent prediction patterns (Fig. 7A, B). In contrast, hemoglobin, serum iron, quantitative CRP, and hematocrit showed relatively low prediction performance in both sexes (R ≤ 0.6)(Fig. 7A). Several biomarkers also showed sex-dependent prediction patterns (Fig. 7A). HbA1c, α2-globulin, free T4, γ-globulin, MCH, Mg, TSH, blood glucose, free T3, triglyceride, total bilirubin, MCV, total cholesterol, and serum amylase were more accurately predicted in males, whereas α1-globulin, γ-GTP, and insulin were more accurately predicted in females. As in cheek mucosa-derived models, the CpG sites selected in the male and female blood-derived models showed limited overlap for most biomarkers, and approximately 20% of the selected CpG sites overlapped with sex-associated CpG sites identified by genome-wide analysis (Fig. 8).

**Figure 6.**
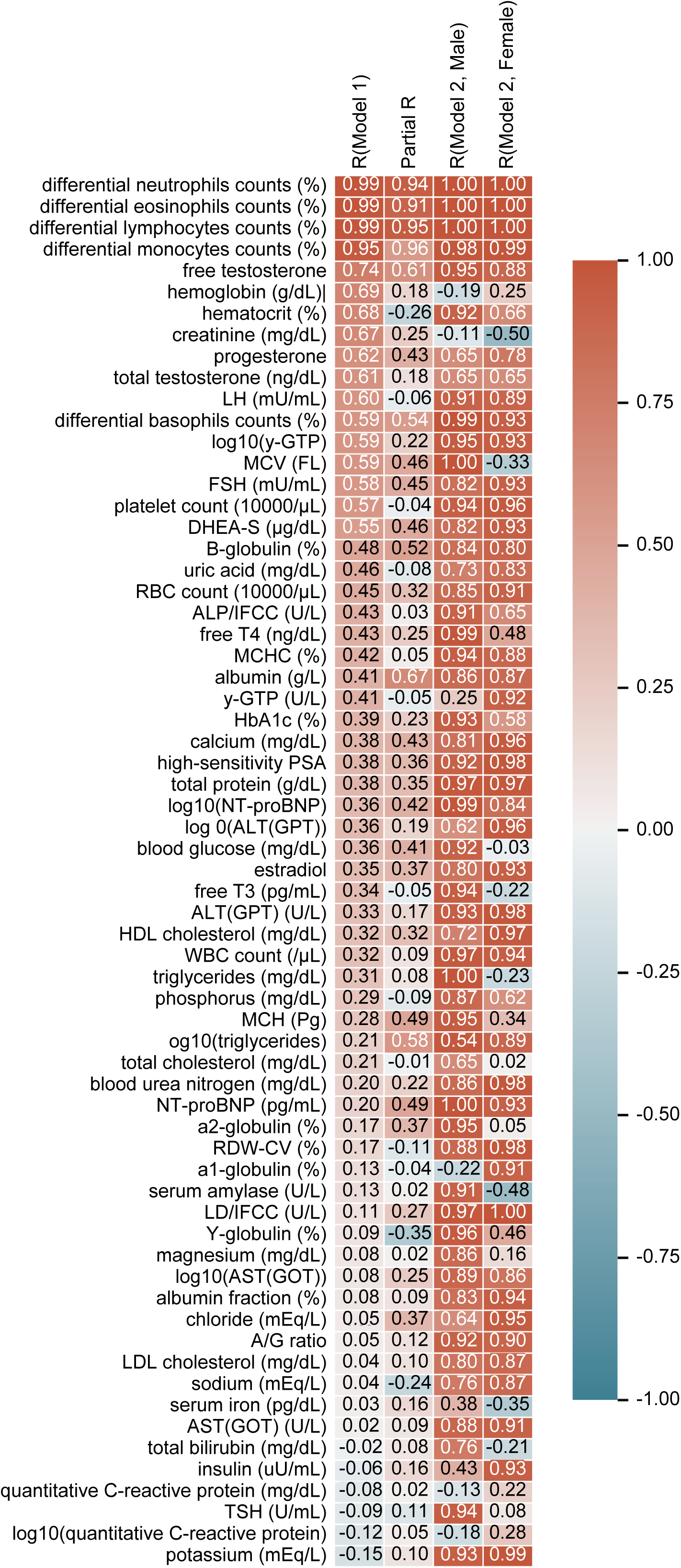
Prediction performance for clinical biomarkers based on blood-derived MSA methylation data. Heatmap showing the correlation coefficients between observed and predicted values for all 59 clinical biomarkers. Model 1 represents prediction performance across all participants and is shown as the Pearson correlation coefficient (R, Model 1). Partial R indicates the sex-adjusted partial correlation coefficient. Model 2 represents sex-stratified prediction performance and is shown separately for males (R, Model 2 Male) and females (R, Model 2 Female).

**Figure 7.**
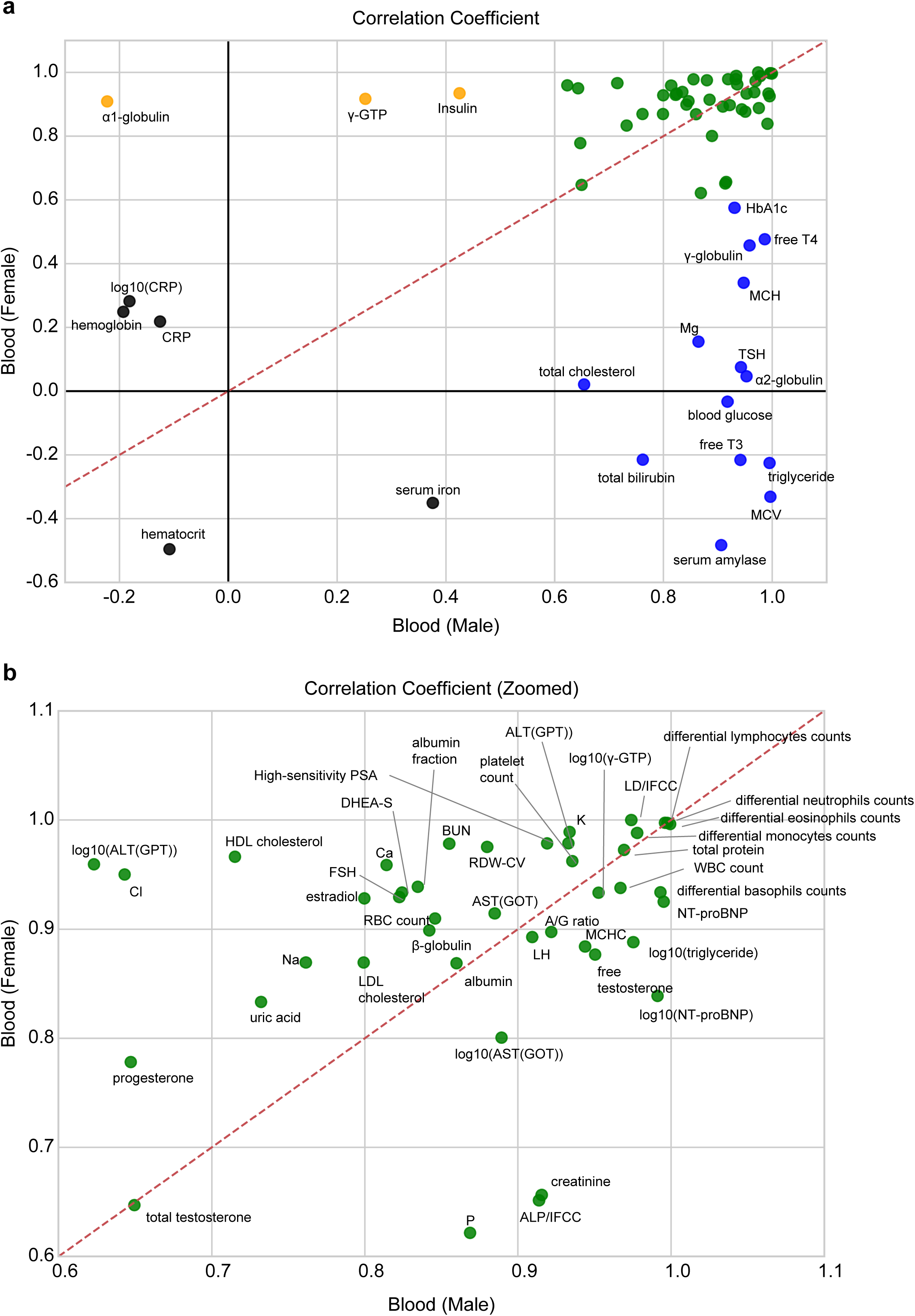
Comparison of biomarker prediction performance between males and females based on blood-derived MSA methylation data. Scatter plot showing the comparison of correlation coefficients obtained from Model 2 for all 59 clinical biomarkers between males and females (A). Panel (B) shows an enlarged view of panel (A). Biomarkers with correlation coefficients ≥ 0.6 in both sexes are shown in green, those with correlation coefficients ≥ 0.6 in males only are shown in blue, and those with correlation coefficients ≥ 0.6 in females only are shown in orange. All other biomarkers are shown in black. The red dashed line indicates the line of identity (y = x).

**Figure 8.**
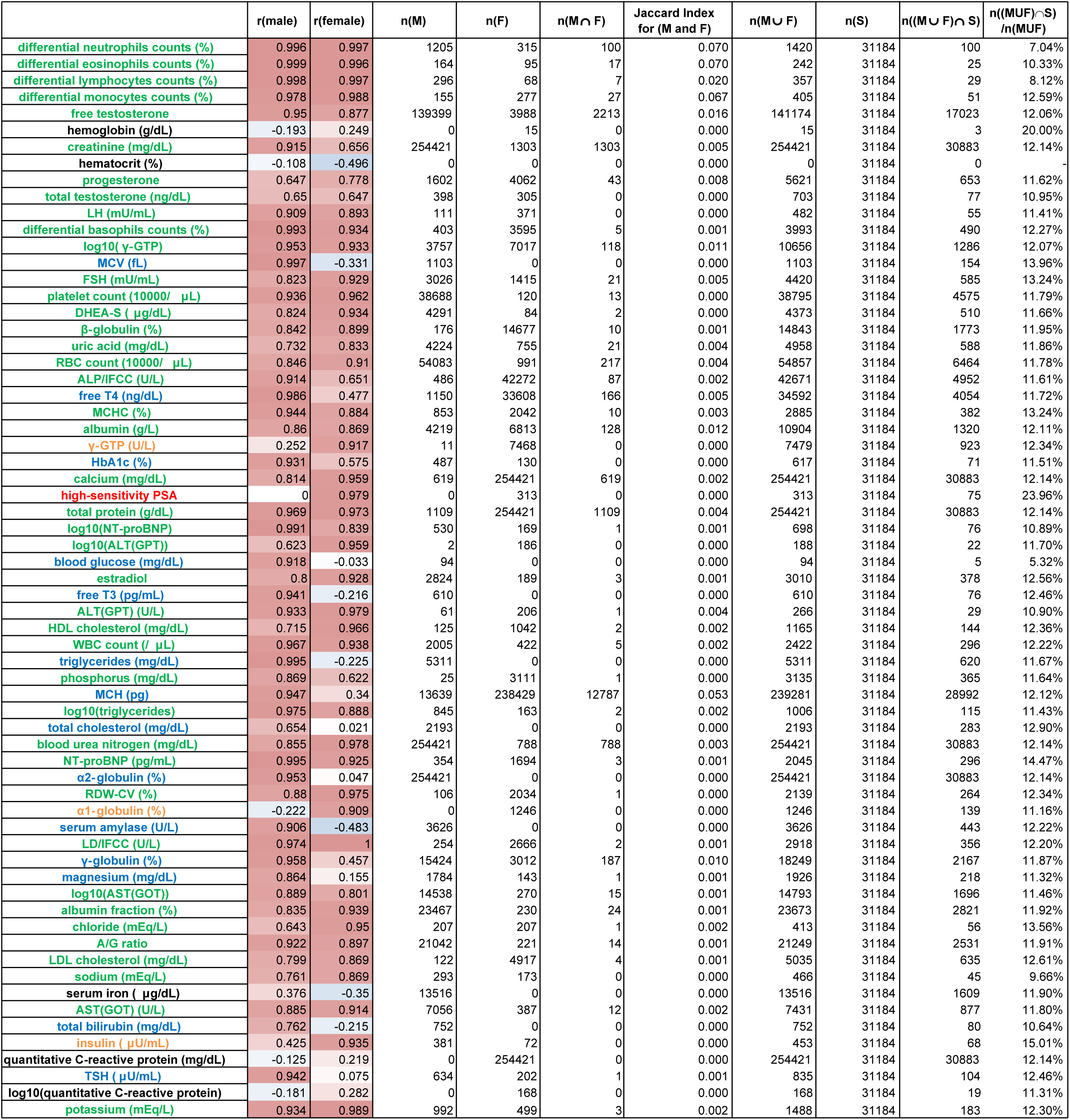
Comparison of CpG features between male and female models based on blood-derived DNA methylation and their overlap with sex-associated CpG sites. For all 59 clinical biomarkers, the prediction performance of Model 2 is shown as correlation coefficients for males (R [Male]) and females (R [Female]), with text colors corresponding to Figure 7. The number of CpG sites selected in male (M) and female (F) models, their overlap (n(M ∩ F)), and the Jaccard index are presented, with values ≥ 0.8 highlighted in yellow. When M ∪ F contained no CpG sites, the Jaccard index was set to 0. Overlap with sex-associated CpG sites (S) is evaluated using the number of CpGs in M ∪ F, the number of overlapping CpGs (n((M ∪ F) ∩ S)), and their proportion. When M ∪ F contained no CpG sites, the value is indicated as a hyphen.

### Comparison of Biomarker Prediction Between Cheek Mucosa-Derived and Blood-Derived MSA Data

Finally, we compared biomarker prediction performance between cheek mucosa-derived and blood-derived MSA methylation data separately in males and females. In males, biomarkers that were relatively well predicted from cheek mucosa-derived data were generally well predicted from blood-derived data, although prediction performance differed for several markers, including quantitative CRP, serum iron, insulin, AST(GOT), Cl, triglycerides, γ-globulin, MCH, β-globulin, total bilirubin, FSH, free T3, NT-proBNP, ALT(GPT), RBC count, uric acid, and ALT (GPT)(Fig. 9A, B). A similar overall pattern was observed in females, with substantial overlap in the set of biomarkers that could be predicted from cheek mucosa-derived and blood-derived methylation data, despite differences in prediction performance for individual markers including free T4, γ-globulin, MCH, quantitative CRP, Mg, TSH, blood glucose, free T3, serum iron, hematocrit, LDL cholesterol, creatine, uric acid, differential basophiles counts, insulin, RBC count, and α1-globulin (Fig. 10A, B). Sex- and tissue-dependent predictability of 59 biomarkers are summarized with functional categories in Fig. 11. We next compared the CpG sites selected in the sex-stratified biomarker prediction models between cheek mucosa-derived and blood-derived MSA data. In both males and females, the overlap between the CpG sites selected in the two tissues was minimal for most biomarkers, as indicated by low Jaccard indices (Fig. 12 for males and Fig. 13 for females). In contrast, most CpG sites selected in the cheek mucosa-derived or blood-derived models overlapped with CpG sites exhibiting significant tissue-associated methylation differences, as defined by genome-wide analysis (Fig. 12 and Fig. 13). Thus, although similar clinical biomarkers could be predicted from cheek mucosa-derived and blood-derived methylation data, the corresponding prediction models were constructed using largely distinct and tissue-associated CpG features.

**Figure 9.**
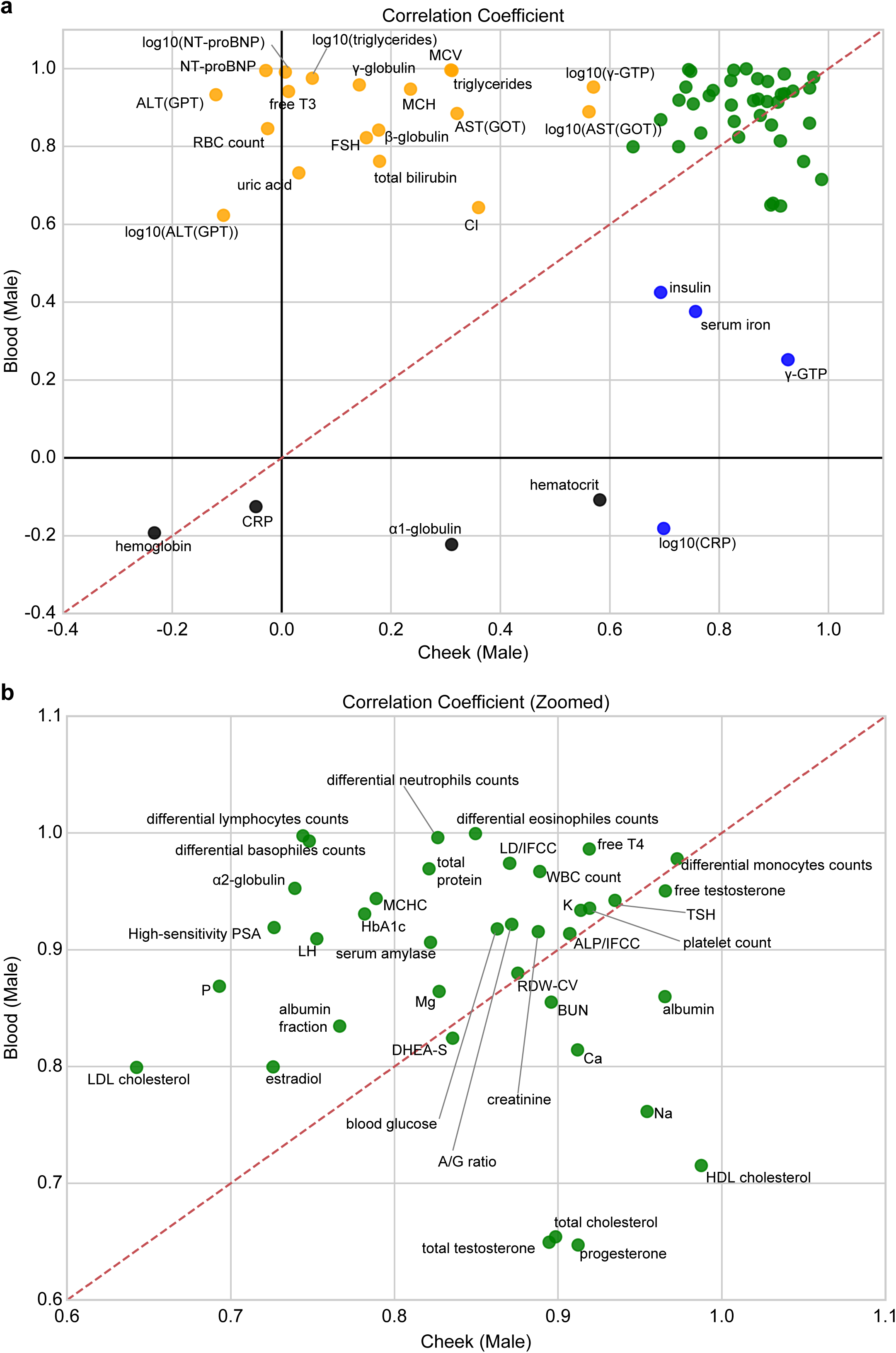
Comparison of biomarker prediction performance between cheek mucosa- and blood-derived MSA methylation data in males. Scatter plot showing the comparison of correlation coefficients obtained from Model 2 for all 59 clinical biomarkers between cheek mucosa-derived and blood-derived MSA methylation data (A). Panel (B) shows an enlarged view of panel (A). Biomarkers with correlation coefficients ≥ 0.6 in both models are shown in green, those with correlation coefficients ≥ 0.6 in the cheek mucosa-derived model only are shown in blue, and those with correlation coefficients ≥ 0.6 in the blood-derived model only are shown in orange. All other biomarkers are shown in black. The red dashed line indicates the line of identity (y = x).

**Figure 10.**
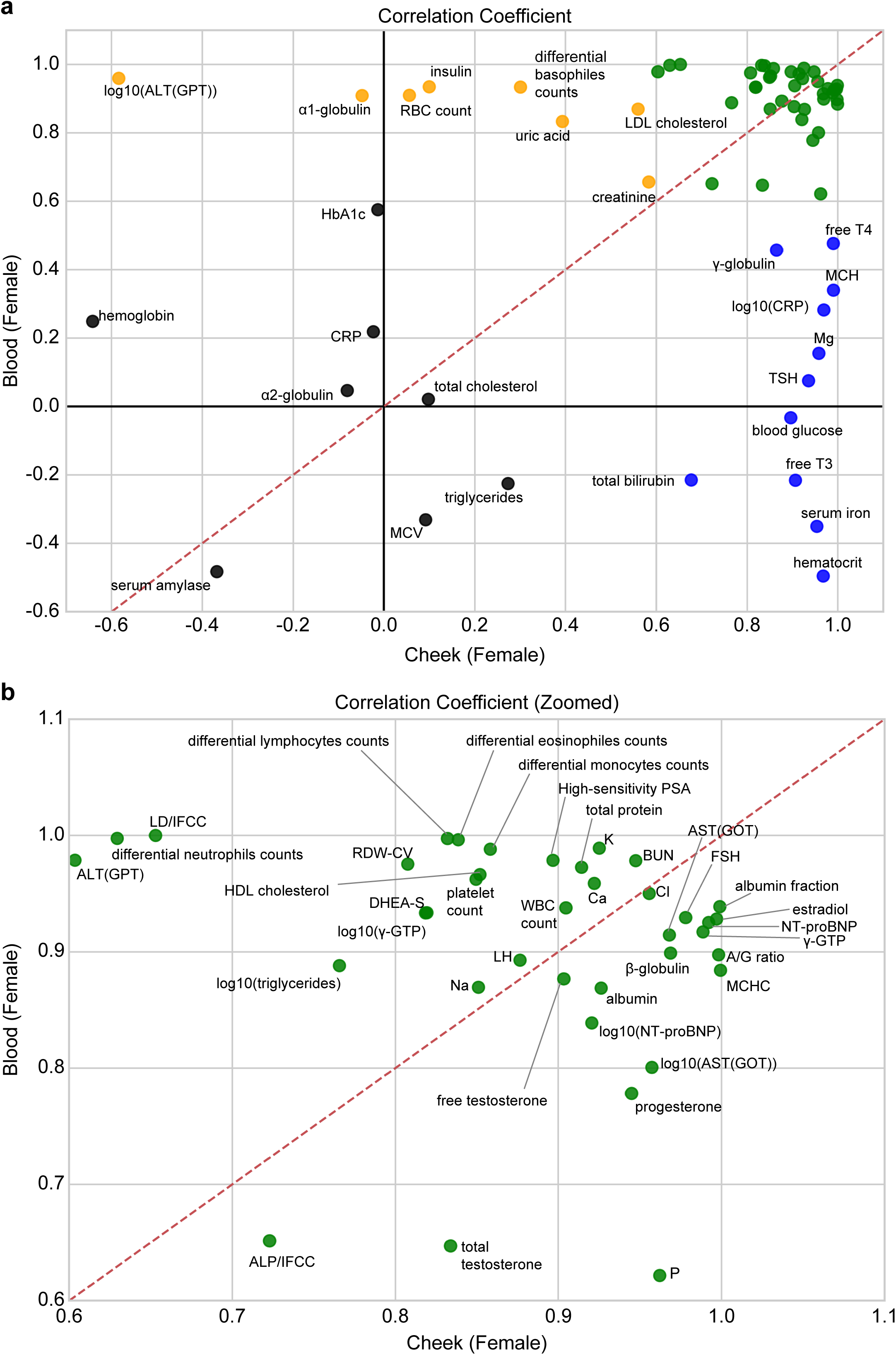
Comparison of biomarker prediction performance between cheek mucosa- and blood-derived MSA methylation data in females. Scatter plot showing the comparison of correlation coefficients obtained from Model 2 for all 59 clinical biomarkers between cheek mucosa-derived and blood-derived MSA methylation data (A). Panel (B) shows an enlarged view of panel (A). Biomarkers with correlation coefficients ≥ 0.6 in both models are shown in green, those with correlation coefficients ≥ 0.6 in the cheek mucosa-derived model only are shown in blue, and those with correlation coefficients ≥ 0.6 in the blood-derived model only are shown in orange. All other biomarkers are shown in black. The red dashed line indicates the line of identity (y = x).

**Figure 11.**
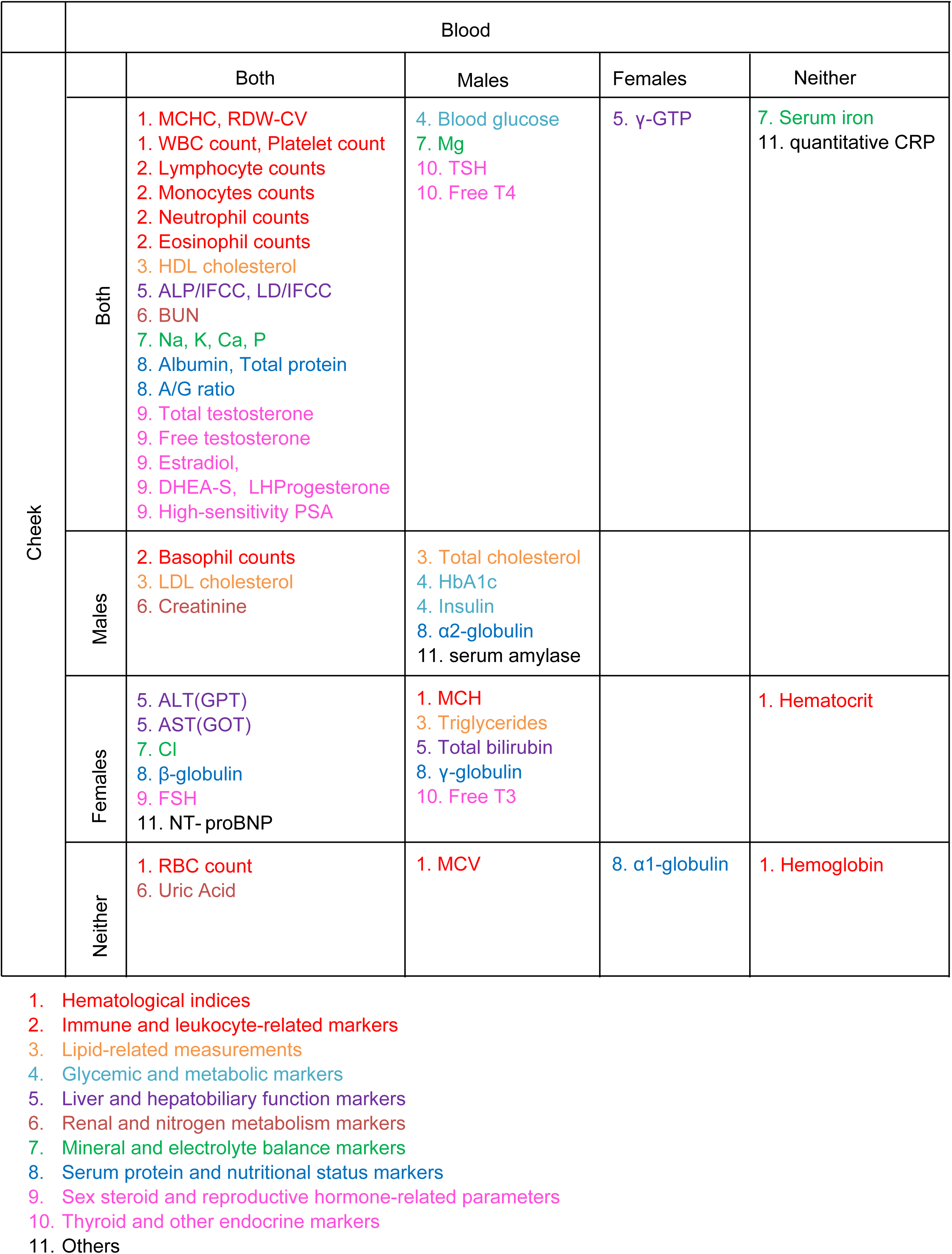
Sex- and tissue-dependent predictability of clinical biomarkers and their functional categories. The predictability of 59 clinical biomarkers is categorized based on Model 2’s performance in cheek mucosa-derived and blood-derived methylation models, each stratified by sex. For each tissue, biomarkers are classified into four categories: Both (correlation coefficient r > 0.6 in both sexes), Males (r > 0.6 in males only), Females (r > 0.6 in females only), and Neither (r < 0.6 in both sexes), resulting in a total of 16 classification patterns. Each biomarker is additionally annotated with a number and color according to its biological functional category.

**Figure 12.**
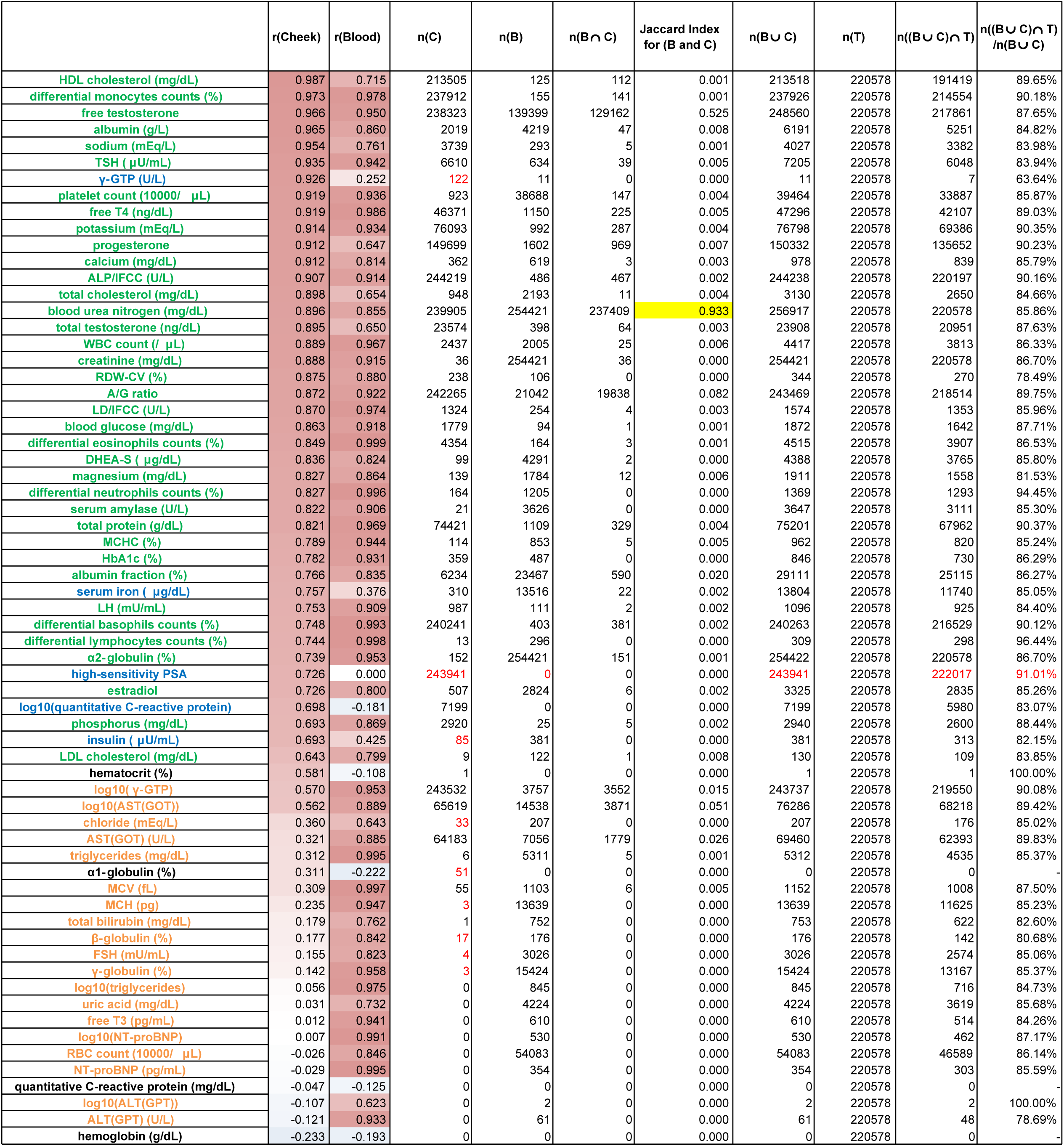
Comparison of CpG features between cheek mucosa-derived and blood-derived models in males and their overlap with tissue-associated CpG sites. For all 59 clinical biomarkers, the prediction performance of Model 2 is shown as correlation coefficients for males (R [Cheek]) and females (R [Blood]), with text colors corresponding to Figure 9. The number of CpG sites selected in cheek mucosa-derived (C) and blood-derived (B) models, their overlap (n(B ∩ C)), and the Jaccard index are presented, with values ≥ 0.8 highlighted in yellow. When B ∪ C contained no CpG sites, the Jaccard index was set to 0. Overlap with tissue-associated CpG sites (T) is evaluated using the number of CpGs in B ∪ C, the number of overlapping CpGs (n((B ∪ C) ∩ T)), and their proportion. When B ∪ C contained no CpG sites, the value is indicated using a hyphen.

**Figure 13.**
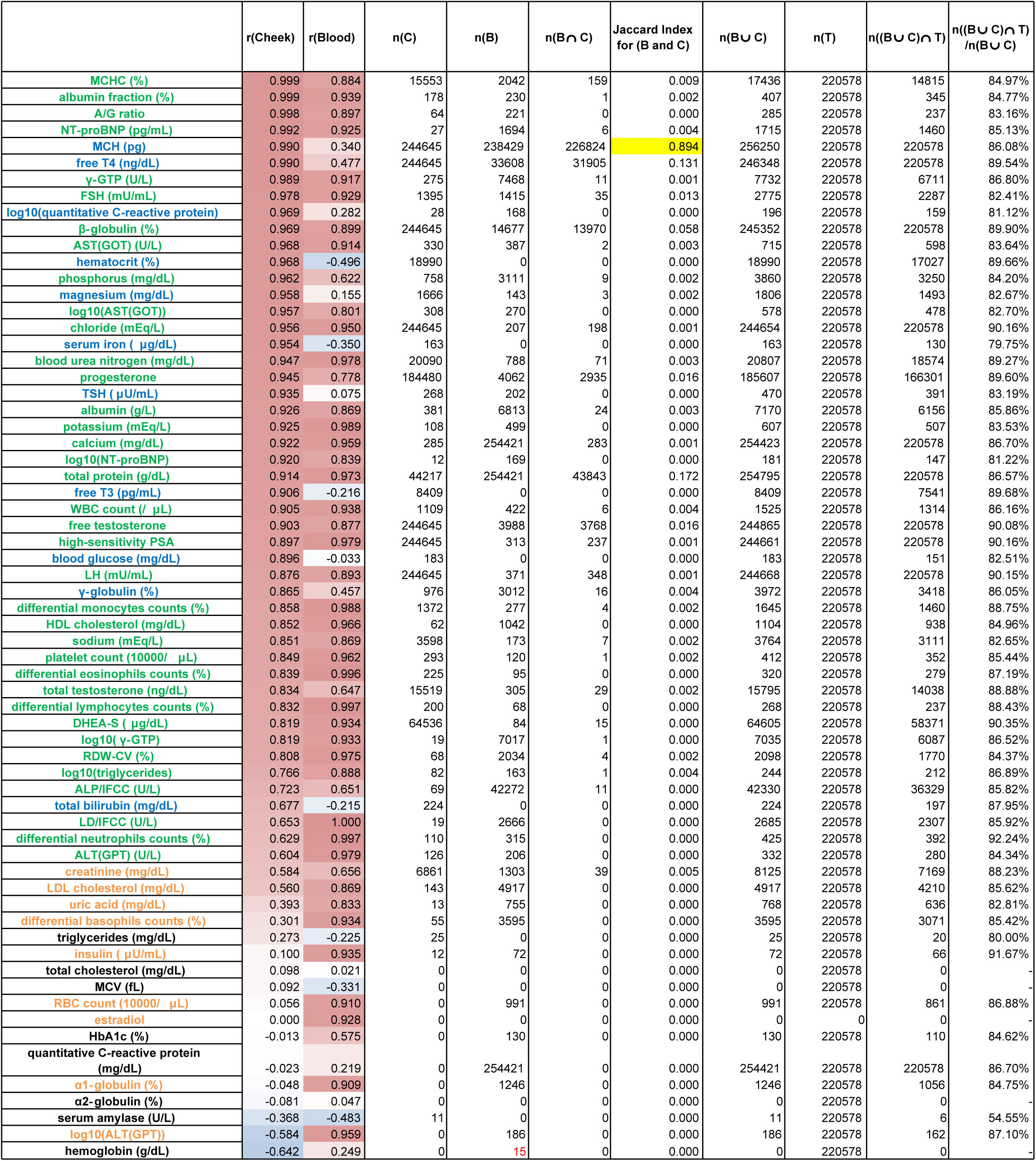
Comparison of CpG features between cheek mucosa-derived and blood-derived models in females and their overlap with tissue-associated CpG sites. For all 59 clinical biomarkers, the prediction performance of Model 2 is shown as correlation coefficients for males (R [Cheek]) and females (R [Blood]), with text colors corresponding to Figure 10. The number of CpG sites selected in cheek mucosa-derived (C) and blood-derived (B) models, their overlap (n(B ∩ C)), and the Jaccard index are presented, with values ≥ 0.8 highlighted in yellow. When B ∪ C contained no CpG sites, the Jaccard index was set to 0. Overlap with tissue-associated CpG sites (T) is evaluated using the number of CpGs in B ∪ C, the number of overlapping CpGs (n((B ∪ C) ∩ T)), and their proportion. When B ∪ C contained no CpG sites, the value is indicated as a hyphen.

## Discussion

This study demonstrated that DNA methylation profiles obtained from cheek mucosa using the MSA could predict chronological age, phenotypic age, and a range of clinically relevant laboratory biomarkers in a Japanese cohort. The two-stage residual learning approach substantially improved prediction performance for phenotypic age, indicating that information captured by established blood-based epigenetic clocks could be effectively leveraged to refine aging-related predictions from a less invasive oral sample and cheek mucosa-derived MSA-specific methylation signals successfully captured residual variance not explained by the clocks. Multiple physiological biomarkers could be predicted from cheek mucosa-derived methylation data, although prediction performance varied across markers and was strongly influenced by sex. Finally, comparative feature analyses revealed that biomarker prediction models often relied on markedly different CpG sets across sexes and tissues, despite targeting similar phenotypes. These findings support the potential utility of cheek mucosa as an accessible substrate for methylation-based assessment of biological aging and physiological state, while highlighting the importance of tissue- and sex-aware model design.

The strong performance of the cheek mucosa-derived models for chronological age prediction is consistent with the notion that age-associated DNA methylation changes include a substantial component shared across tissues. Although many methylation changes are tissue-specific, a subset of age-related CpG alterations is reproducibly observed across multiple cell and tissue types, thereby supporting the robustness of epigenetic age estimation beyond blood alone [3, 10]. Consistent with this interpretation, 150 CpG sites were shared between the blood-derived MSA-based one-stage chronological age model (207 CpGs) and the cheek mucosa-derived MSA-based one-stage chronological age model (289 CpGs). Our findings, therefore, suggest that cheek mucosa retains practically sufficient age-related methylation structure to support accurate prediction of chronological age even when measured using a reduced-coverage platform such as the MSA. By contrast, the comparatively weaker concordance with blood-derived PhenoAge and the greater improvement achieved by the two-stage approach for phenotypic age prediction imply that clinically relevant aging phenotypes may depend more strongly on tissue-contextual and model-specific methylation information than chronological aging alone. This contrast was reflected in the selected model features, as only four CpG sites were shared between the blood-derived MSA-based one-stage phenotypic age model (364 CpGs) and the cheek mucosa-derived MSA-based one-stage phenotypic age model (174 CpGs). In this context, the two-stage approach may have been effective because it allowed broad aging-related information captured by established blood-based clocks to be retained while simultaneously incorporating tissue- and platform-specific methylation features from cheek mucosa. This interpretation is consistent with the idea that phenotypic age reflects a more complex composite of systemic physiological dysregulation than chronological age, consequently requiring both shared and tissue-specific epigenetic information for improved prediction [2].

The marked improvement in biomarker prediction performance after sex stratification suggests that a substantial proportion of the methylation signal relevant to physiological traits is sex-dependent and may be obscured in models trained across both sexes. This interpretation is biologically plausible, given the well-established sex differences in many laboratory biomarkers and the widespread sex-associated variation in DNA methylation [24, 25]. The limited overlap in CpG features selected by male and female models further supports the idea that similar biomarkers may be encoded through partially distinct epigenetic architectures in each sex. Simultaneously, the observation that many biomarkers predictable from blood-derived methylation data were also predictable from cheek mucosa-derived methylation data suggests that some clinically relevant physiological states are reflected in methylation patterns shared across tissues. This appeared to be particularly true for biomarkers related to hematologic, immune, electrolyte, and sex steroid functions, whereas glycemic, lipid-related, hepatobiliary, thyroid-related, and renal markers showed greater sex-and tissue-dependence. Such differences may reflect variation in how strongly each physiological domain is embedded in peripheral methylation states, with some traits more robustly represented across tissues and others requiring more context-specific epigenetic information [3, 10].

One particularly notable finding was that quantitative CRP, which has not consistently shown strong predictability from blood-derived DNA methylation data alone [2, 15], was predicted with relatively high accuracy from cheek mucosa-derived methylation profiles in the sex-stratified analyses. This result may be clinically relevant because CRP is a widely used marker of low-grade systemic inflammation and is closely linked to cardiometabolic risk, frailty, and biological aging [15, 26]. Oral mucosal methylation may reflect aspects of chronic inflammatory exposure or immune–environment interactions that are not fully represented by blood cell methylation alone [27, 28], particularly in a tissue that is continuously exposed to microbial, dietary, and environmental stimuli [29, 30]. This raises the possibility that cheek mucosa-derived methylation profiling can provide a non-invasive readout of chronic inflammatory burden. The predictability of biomarkers related to insulin metabolism, thyroid function, and sex hormone regulation may also have important translational implications. Markers such as insulin, free T3, free T4, testosterone, estradiol, LZ, FSH, and progesterone are tightly linked to endocrine and metabolic homeostasis, and their dysregulation is associated with a broad range of age-related and chronic disorders [31–35]. The ability to predict these biomarkers from cheek mucosa- or blood-derived methylation data suggests that epigenetic patterns in peripheral tissues may encode downstream physiological consequences of hormonal and metabolic signaling, even when the source tissues are not classical endocrine organs. This observation is clinically relevant because it raises the possibility that methylation-based models may eventually complement conventional blood testing to assess systemic physiological state, particularly in longitudinal or screening-oriented settings where repeated, less invasive sampling is desirable.

An additional key finding of this study was that biomarker prediction models constructed separately by sex or tissue relied on largely distinct sets of CpG features, despite targeting similar physiological outcomes. This pattern was consistently observed across both sex-stratified and cross-tissue comparisons, where minimal overlap in selected CpG sites was accompanied by substantial differences in feature composition. Many biomarkers remained predictable across sexes and tissues, indicating that similar phenotypic information could be encoded through multiple, partially independent epigenetic configurations. One possible explanation for this apparent discrepancy is that DNA methylation does not represent a single deterministic mapping to physiological traits but instead reflects a high-dimensional and context-dependent regulatory landscape. In this framework, different subsets of CpG sites may capture correlated downstream effects of shared upstream biological processes, such as hormonal signaling, immune activation, or metabolic regulation, even if the specific loci involved differ by sex or tissue. Differences in cellular composition, local microenvironment, and exposure history between tissues may further shape the methylation features selected by predictive models. Thus, the observed divergence in CpG feature sets likely reflects both biological heterogeneity and model flexibility, rather than reflecting inconsistency in the underlying physiological signals.

Limitations of this study warrant further consideration. Since the analyses were conducted in a single Japanese cohort, the generalizability of the developed models to other populations, age distributions, and clinical backgrounds remains to be examined. Specifically, although we used held-out test sets, external validation in fully independent cohorts is essential to establish the generalizability of both the aging-related and biomarker prediction models. Despite these limitations, our findings highlight the potential of cheek mucosa as a practical and biologically informative substrate for methylation-based phenotyping. Compared with blood collection, cheek sampling is less invasive, readily repeatable, and more easily deployable in large-scale or longitudinal settings. If validated in independent and more diverse cohorts, cheek mucosa-derived methylation profiling may provide a useful approach for non-invasive prediction of biological aging, systemic physiological state, and clinically relevant biomarker patterns. These features may be particularly valuable for future screening, monitoring, and personalized health applications.

## Conclusion

In conclusion, DNA methylation profiles derived from cheek mucosa using the Illumina Infinium MSA enabled the prediction of chronological age, phenotypic age, and multiple clinically relevant laboratory biomarkers in this Japanese cohort. The two-stage residual learning approach was particularly effective in improving the prediction of phenotypic age, suggesting that integrating information from established blood-based epigenetic clocks with cheek mucosa-derived methylation features may enhance the estimation of complex aging-related phenotypes. The predictability of inflammatory, metabolic, endocrine, and hormonal biomarkers from cheek mucosa-derived methylation data highlights the broader physiological information contained in oral epithelial methylation profiles. The marked differences in selected CpG features across sexes and tissues further indicate that biologically informed, context-aware modeling strategies are important for methylation-based phenotyping. Together, these findings support the potential of cheek mucosa as a less invasive alternative for the assessment of biological aging and systemic physiological state.

## Additional Files

Additional File 1. Proportion of missing values for each of the 59 clinical laboratory measurements.

Additional File 2. Imputed values for all 59 clinical laboratory measurements for 186 participants.

Additional File 3. Estimated coefficients in the one-stage model.

Additional File 4. Estimated coefficients in the two-stage model.

Additional File 5. MAE and RMSE between observed and predicted values in Models 1 and 2 based on cheek mucosa- and blood-derived MSA data.

Additional File 6. Estimated coefficients in Model 1 for biomarkers based on cheek mucosa-derived MSA data.

Additional File 7. Estimated coefficients in Model 2 (Males) for biomarkers based on cheek mucosa-derived MSA data.

Additional File 8. Estimated coefficients in Model 2 (Females) for biomarkers based on cheek mucosa-derived MSA data.

Additional File 9. Estimated coefficients in Model 1 for biomarkers based on blood-derived MSA data.

Additional File 10. Estimated coefficients in Model 2 (Males) for biomarkers based on blood-derived MSA data.

Additional File 11. Estimated coefficients in Model 2 (Females) for biomarkers based on blood-derived MSA data.

## List of Abbreviations

MSA: Methylation screening array
MCV: mean cell volume
RDW-CV: red cell distribution width-coefficient of variation
ALP: alkaline phosphatase
WBC: white blood cell
J-PhenoAge: Japanese PhenoAge
ALP/IFCC: International Federation of Clinical Chemistry and Laboratory Medicine
ALT: alanine aminotransferase
GPT: glutamic pyruvate transaminase
AST: aspartate aminotransferase
DHEA-S: dehydroepiandrosterone sulfate
FSH: follicle-stimulating hormone
HDL: high-density lipoprotein
Hb: hemoglobin
LD/IFCC: lactate dehydrogenase based on IFCC
LH: luteinizing hormone
MCH: mean corpuscular Hb
NT-proBNP: N-terminal pro-brain natriuretic peptide
RBC: red blood cell
γ-GTP: γ-glutamyl transpeptidase
free T3: free triiodothyronine
free T4: free thyroxine
A/G ratio: albumin/globulin ratio
PSA: prostate-specific antigen
MS: mortality score
MAE: mean absolute error
RMSE: root mean squared error

## Declarations

### Ethics approval and consent to participate

A total of 186 samples were collected at Y’s Science Clinic Hiroo (Minato-ku, Tokyo, Japan) as part of the project titled “Evaluation of Biological Age Based on DNA Methylation and Its Clinical Significance,” reviewed and approved by the Shiba Palace Clinic’s Institutional Review Board. Written informed consent was obtained from all participants.

### Consent for publication

Not applicable.

### Availability of data and materials

The imputed values for all clinical laboratory measurements are provided in Additional file 2. The original values are available upon reasonable request. Estimated coefficients for each model are provided in Additional files 3, 4, and 6. MAE and RMSE are summarized in Additional file 5. All other data supporting the findings of this study are available from the corresponding author upon reasonable request.

### Author Contributions

Tatsuma Shoji (TS) performed data analysis, generated all figures and tables, and wrote the manuscript draft. Yui Tomo (YT) provided statistical guidance, contributed to the design of the analytical framework, and reviewed and edited the manuscript draft. Ryo Nakaki (RN) collected the data and led the overall direction and management of the project.

## Acknowledgements

We extend our deepest gratitude to Sawako Hibino, whose efforts in coordinating and collecting blood samples formed the basis of the dataset used in this study. We thank Editage (www.editage.com) for providing assistance in English language editing. The contributions of these individuals were indispensable for the successful execution of this study.

## References

1. Horvath S, Raj K. DNA methylation-based biomarkers and the epigenetic clock theory of ageing. Nat Rev Genet. 2018;19:371–84. 10.1038/s41576-018-0004-3

2. Levine ME, Lu AT, Quach A, Chen BH, Assimes TL, Bandinelli S, et al. An epigenetic biomarker of aging for lifespan and healthspan. Aging. 2018;10:573–91. 10.18632/aging.101414

3. Horvath S. DNA methylation age of human tissues and cell types. Genome Biol. 2013;14:R115. 10.1186/gb-2013-14-10-r115

4. Belsky DW, Caspi A, Houts R, Cohen HJ, Corcoran DL, Danese A, et al. Quantification of biological aging in young adults. Proc Natl Acad Sci USA. 2015;112:E4104–E10. 10.1073/pnas.1506264112

5. Belsky DW, Caspi A, Arseneault L, Baccarelli A, Corcoran DL, Gao X, et al. Quantification of the pace of biological aging in humans through a blood test, the DunedinPoAm DNA methylation algorithm. eLife. 2020;9:e54870. 10.7554/eLife.54870

6. Liu Z, Kuo PL, Horvath S, Crimmins E, Ferrucci L, Levine M. A new aging measure captures morbidity and mortality risk across diverse subpopulations from NHANES IV: A cohort study. PLOS Med. 2018;15:e1002718. 10.1371/journal.pmed.1002718

7. Hannum G, Guinney J, Zhao L, Zhang L, Hughes G, Sadda S, et al. Genome-wide methylation profiles reveal quantitative views of human aging rates. Mol Cell. 2013;49:359–67. 10.1016/j.molcel.2012.10.016

8. Lu AT, Quach A, Wilson JG, Reiner AP, Aviv A, Raj K, et al. DNA methylation GrimAge strongly predicts lifespan and healthspan. Aging. 2019;11:303–27. 10.18632/aging.101684

9. Lowe R, Gemma C, Beyan H, Hawa MI, Bazeos A, Leslie RD, et al. Buccals are likely to be a more informative surrogate tissue than blood for epigenome-wide association studies. Epigenetics. 2013;8:445–54. 10.4161/epi.24362

10. Slieker RC, Relton CL, Gaunt TR, Slagboom PE, Heijmans BT. Age-related DNA methylation changes are tissue-specific with ELOVL2 promoter methylation as exception. Epigenet Chromatin. 2018;11:25. 10.1186/s13072-018-0191-3

11. McEwen LM, O’Donnell KJ, McGill MG, Edgar RD, Jones MJ, MacIsaac JL, et al. The PedBE clock accurately estimates DNA methylation age in pediatric buccal cells. Proc Natl Acad Sci USA. 2020;117:23329–35. 10.1073/pnas.1820843116

12. Eipel M, Mayer F, Arent T, Ferreira MRP, Birkhofer C, Gerstenmaier U, et al. Epigenetic age predictions based on buccal swabs are more precise in combination with cell type-specific DNA methylation signatures. Aging. 2016;8:1034–48. 10.18632/aging.100972

13. Johnson AA, Torosin NS, Shokhirev MN, Cuellar TL. A set of common buccal CpGs that predict epigenetic age and associate with lifespan-regulating genes. iScience. 2022;25:105304. 10.1016/j.isci.2022.105304

14. Shokhirev MN, Torosin NS, Kramer DJ, Johnson AA, Cuellar TL. CheekAge: a next-generation buccal epigenetic aging clock associated with lifestyle and health. GeroScience. 2024;46:3429–43. 10.1007/s11357-024-01094-3

15. Ligthart S, Marzi C, Aslibekyan S, Mendelson MM, Conneely KN, Tanaka T, et al. DNA methylation signatures of chronic low-grade inflammation are associated with complex diseases. Genome Biol. 2016;17:255. 10.1186/s13059-016-1119-5

16. Tomo Y, Nakaki R. Transfer Elastic Net for developing epigenetic clocks for the Japanese population. Mathematics. 2024;12:2716. 10.3390/math12172716

17. Shoji T, Tomo Y, Nakaki R. Prediction of biological age and blood biomarkers from DNA methylation profiles measured by the Methylation Screening Array: development and validation of models on Japanese data. bioRxiv. 2026. 10.64898/2026.02.06.703638

18. Geurts P, Ernst D, Wehenkel L. Extremely randomized trees. Mach Learn. 2006;63:3–42. 10.1007/s10994-006-6226-1

19. Aryee MJ, Jaffe AE, Corrada-Bravo H, Ladd-Acosta C, Feinberg AP, Hansen KD, et al. Minfi: a flexible and comprehensive Bioconductor package for the analysis of Infinium DNA methylation microarrays. Bioinformatics. 2014;30:1363–9. 10.1093/bioinformatics/btu049

20. Zou H, Hastie T. Regularization and variable selection via the elastic net. J R Stat Soc B. 2005;67:301–20. 10.1111/j.1467-9868.2005.00503.x

21. Kuzborskij I, Orabona F. Stability and hypothesis transfer learning. In: Proceedings of the 30th international conference on machine learning (ICML 2013). Proceedings of Machine Learning Research; 2013;p. 942–50

22. Lu AT, Binder AM, Zhang J, Yan Q, Reiner AP, Cox SR, et al. DNA methylation GrimAge version 2. Aging. 2022;13:4781–800. 10.18632/aging.204434

23. Zheng G, Liu C, Wei H, Jenkins P, Chen C, Wen T, et al. Knowledge-based residual learning. In: Proceedings of the thirtieth international joint conference on artificial intelligence. International Joint Conferences on Artificial Intelligence Organization; 2021;p. 1653–9. 10.24963/ijcai.2021/228

24. Singmann P, Shem-Tov D, Wahl S, Grallert H, Fiorito G, Shin SY, et al. Characterization of whole-genome autosomal differences of DNA methylation between men and women. Epigenet Chromatin. 2015;8:15. 10.1186/s13072-015-0035-3

25. Yusipov I, Bacalini MG, Kalyakulina A, Krivonosov M, Pirazzini C, Gensous N, et al. Age-related DNA methylation changes are sex-specific: a comprehensive assessment. Aging. 2020;12:24057–80. 10.18632/aging.202251

26. Franceschi C, Garagnani P, Parini P, Giuliani C, Santoro A. Inflammaging: a new immune-metabolic viewpoint for age-related diseases. Nat Rev Endocrinol. 2018;14:576–90. 10.1038/s41574-018-0059-4

27. Bonder MJ, Kasela S, Kals M, Tamm R, Lokk K, Barragan I, et al. Genetic and epigenetic regulation of gene expression in fetal and adult human livers. BMC Genomics. 2014;15:860. 10.1186/1471-2164-15-860

28. Smith AK, Kilaru V, Klengel T, Mercer KB, Bradley B, Conneely KN, et al. DNA extracted from saliva for methylation studies of psychiatric traits: evidence tissue specificity and relatedness to brain. Am J Med Genet B. 2015;168:36–44. 10.1002/ajmg.b.32278

29. Gursoy UK, Könönen E. Understanding the roles of gingival beta-defensins. J Oral Microbiol. 2012;4:15127. 10.3402/jom.v4i0.15127

30. Moutsopoulos NM, Konkel JE. Tissue-specific immunity at the oral mucosal barrier. Trends Immunol. 2018;39:276–87. 10.1016/j.it.2017.08.005

31. Duntas LH, Brenta G. The effect of thyroid disorders on lipid levels and metabolism. Med Clin North Am. 2012;96:269–81. 10.1016/j.mcna.2012.01.012

32. Kosmas CE, Bousvarou MD, Kostara CE, Papakonstantinou EJ, Salamou E, Guzman E. Insulin resistance and cardiovascular disease. J Int Med Res. 2023;51. 10.1177/03000605231164548

33. Maggio M, Lauretani F, Ceda GP. Sex hormones and sarcopenia in older persons. Curr Opin Clin Nutr Metab Care. 2013;16:1. 10.1097/MCO.0b013e32835b6044

34. Allan C. Sex steroids and glucose metabolism. Asian J Androl. 2014;16:232. 10.4103/1008-682X.122589

35. Rochira V, Antonio L, Vanderschueren D. EAA clinical guideline on management of bone health in the andrological outpatient clinic. Andrology. 2018;6:272–85. 10.1111/andr.12470

